# The gut microbiota limits Th2 immunity to *Heligmosomoides polygyrus bakeri* infection

**DOI:** 10.1101/2020.01.30.927111

**Authors:** Gabriel A. Russell, Cynthia Faubert, Elena F. Verdu, Irah L. King

## Abstract

Helminth-induced alterations to the gut microbiota have been shown to affect immune responses at local and peripheral sites. Studies examining helminth-microbiota interactions, however, have been limited due to the practical constraints of performing germ-free experiments with parasites that thrive in microbial-rich conditions to complete their development. The infectious (L3) larvae of the murine helminth *Heligmosomoides polygyrus bakeri* (*Hpb*), for example, are normally reared using a fecal-culture method and therefore are inherently unsuitable for germ-free studies *in vivo*. Herein, we detail an adapted methodology for rearing effectively germ-free *Hpb* larvae that are able to maintain the axenic status of a germ-free host during infection. We validate that these larvae do not display any fitness defects relative to fecal-grown larvae and evoke a comparable immune response *in vivo*. Characterization of axenic *Hpb* infection reveals that the commensal microbiota play a multifaceted role during infection - curbing the anti-*Hpb* Th2 response and directing the resolution of tissue granulomas, while simultaneously promoting parasite fitness. Overall these data demonstrate a mutualistic relationship between commensal microbes, enteric helminths and the infected host.

## INTRODUCTION

Intestinal helminths are ubiquitous pathogens that infect almost two billion people worldwide^1, 2^. While they seldom kill the host, helminths cause a variety of damaging morbidities including anemia, diarrhea, intestinal obstruction, and cognitive impairment^1^. As parasites of the gastrointestinal tract, helminths co-exist alongside the gut microbiota and, in recent years, helminth-microbiota interactions have become a question of rising importance. Although several studies have demonstrated that helminth infection leads to alterations of the composition and function of gut microbes^3, 4, 5, 6^, there remain many unanswered questions regarding the role of the microbiota during helminth infection.

This current knowledge gap is largely due to the difficulty of performing *in vivo* germ-free studies of enteric helminth infection. Such experiments are complicated by the confounding microbiota of many helminth species. For example, *Heligmosomoides polygyrus bakeri* (*Hpb*) - a natural murine parasitic roundworm and established model of chronic helminth infection - require microbe-rich feces to develop from eggs into infectious L3 larvae^7, 8^. Consequently, *Hpb* larvae harbor a wide range of commensals that would contaminate a germ-free host. This inherent incompatibility between important model helminths such as *Hpb* and germ-free *in vivo* studies has greatly limited the scope with which helminth-microbiota interactions are examined.

A recent publication examining the impact of helminth-microbiota interactions on allergic airway inflammation described the use of germ-free *Hpb*^5^. A full methodology for rearing larvae in such a manner, however, was not provided. Additionally, the infectivity and fitness of the resultant larvae, as well as their ability to evoke an immune response in both specific pathogen-free (SPF) and germ-free hosts was not tested. In the current study, we detail an adapted methodology for generating germ-free L3 *Hpb*. We validate that these larvae exhibit robust infectivity and fecundity *in vivo* and elicit a comparable cellular and humoral immune response to fecal-grown *Hpb* in SPF mice. We further demonstrate that axenic infection of germ-free mice with *Hpb* maintains host sterility and results in an enhanced Th2 response, exacerbated intestinal inflammation and compromised parasite fitness. Collectively, our data demonstrate that the commensal gut microbiota benefits both the host and parasite through limiting inflammation and promoting helminth fitness, respectively.

## RESULTS

### A bacterial monoculture and nematode growth media are sufficient to generate culture-sterile L3 *Hpb* larvae

In a laboratory setting, infective (L3) stage *Hpb* are normally grown using a copra-culture method, with very little dissection of growth parameters. In short, the feces of infected mice (containing *Hpb* eggs) are cultured in a moist, dark environment for several days^9^. Therein, the eggs hatch and develop through two free-living stages of larvae (L1 and L2), up to the infective L3 stage^7^. While few details are known about this process, we suspected the major component of feces necessary for *Hpb* growth from egg to L3 to be bacteria. To test this possibility, antibiotic-treated *Hpb* eggs were incubated with a monoculture of *Escehrischia coli (E. coli)* HA107 on nematode growth media (NGM) for 5 days, in an otherwise sterile environment (Figure 1A). HA107 is a previously described auxotroph for m-diaminopimelic acid (m-DAP) and D-alanine (D-ala)^10^, two metabolites not produced by mice. As such, this bacterial strain is unable to replicate or persist *in vitro* or *in vivo* without exogenous supplementation^10^.

**Figure 1.**
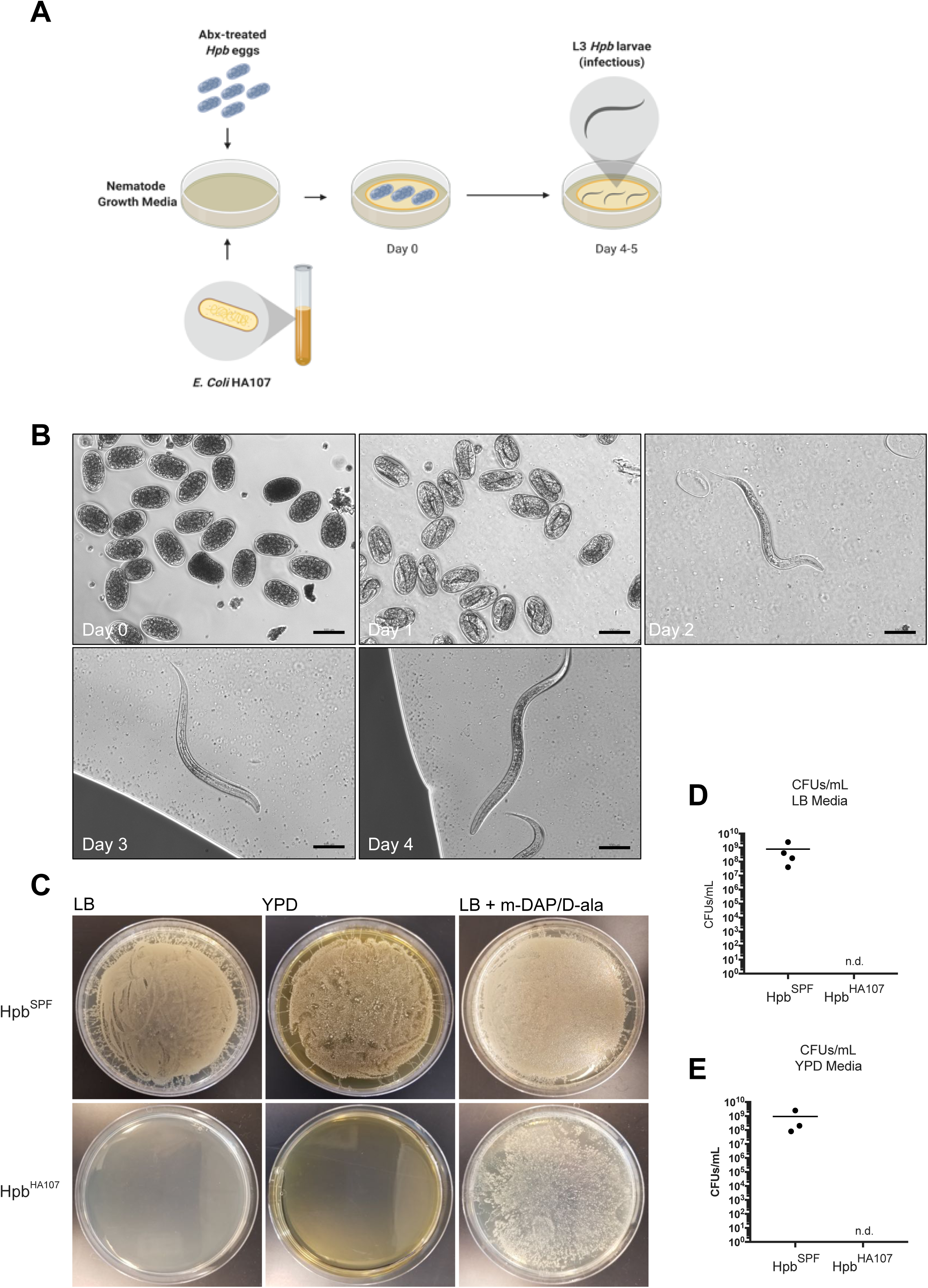
A bacterial monoculture and nematode growth media are sufficient to generate culture-sterile L3 *Hpb* larvae. (A-E) *Hpb* eggs were isolated, sterilized, and incubated with a monoculture of *E. coli* HA107 on Nematode Growth Media in order to generate effectively germ-free L3 stage larvae. (A) Schematic of experimental plan used to generate germ-free *Hpb* larvae. (B) 10X Brightfield microscopy of *Hpb* growth progression in HA107 monoculture. (C) Culture growth from 200 fecal-grown (*Hpb*^SPF^) or HA107-grown (*Hpb*^HA107^) larvae on LB or YPD media. (D-E) Quantification of CFUs resultant from larval culturing. D-ala = D-alanine, m-DAP = m-Diaminopimelic acid, n.d. = non-detectable. Scale bars denote 100µm.

As early as 4 days post-culturing, *Hpb* eggs were found to have successfully reached the L3 larval stage (Figure 1B), as signified by the formation of a cuticle and distinctive movement pattern (not shown). Aerobic cultures of these HA107-grown *Hpb* on both LB and YPD media revealed a complete lack of bacterial or fungal contamination, compared to fecal-grown *Hpb*, effectively rendering the monoculture-grown *Hpb* larvae sterile (Figures 1C-E).

### HA107-grown *Hpb* display comparable infectivity and fecundity and evoke a comparable immune response to fecal-grown parasites

Reduced larval fitness can result in impaired infectivity. Therefore, we assessed adult worm burden of mice infected with fecal-grown *Hpb* (*Hpb*^SPF^) and HA107-reared *Hpb* (*Hpb*^HA107^) at 2 and 4 weeks post-infection. Across both time points, both groups exhibited comparable adult worm burdens (Figure 2A). These results were corroborated by worm fecundity – a measure of parasite fitness (Figure 2B).

**Figure 2.**
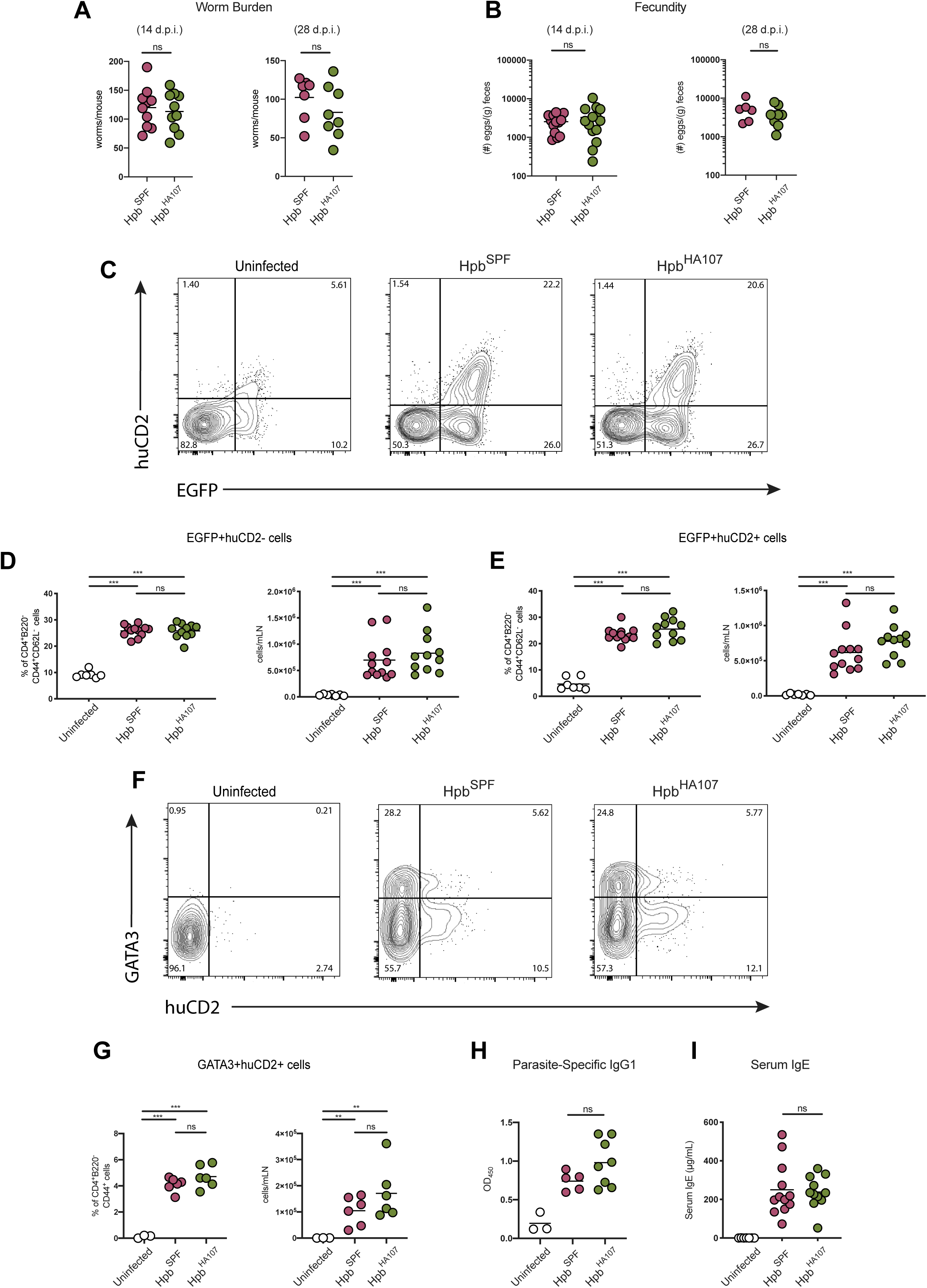
HA107-grown *Hpb* display comparable infectivity, fecundity, and evoke a comparable immune response to fecal-grown parasites. (A-I) 6-12 week-old 4get/KN2 mice were infected with 200 L3 *Hpb*, either HA107-reared (*Hpb*^HA107^) or fecal-grown (*Hpb*^SPF^). (A) Adult worm burden, and (B) fecundity, at 14 and 28 days post infection is shown. (C-G) 14 days post-infection, mesenteric lymph nodes were analyzed by flow cytometry. (C-E) Representative plots, cell numbers, and percentages of live CD4^+^B220^-^CD44^+^CD62L^-^EGFP^+^huCD2^-/+^ cells are shown. (F-G) Representative plots, cell numbers, and percentages of live CD4^+^B220^-^ CD44^+^GATA3^+^huCD2^+^ cells are shown. At 14 days post-infection, serum was extracted and ELISAs for parasite-specific IgG1 (H) and total IgE (I) are shown. Statistical analysis was performed using a t test with Welch’s correction. * indicates p < 0.05, ** indicates p < 0.01, *** indicates p < 0.001, ns = non-significant.

We next sought to validate whether *Hpb*^HA107^ could elicit a comparable immune response to *Hpb*^SPF^ *in vivo.* The anti-*Hpb* cellular immune response is characterized by a robust expansion of CD4^+^ T cells, including T follicular helper (Tfh) and Th2 cells^11^. To track these cells during infection, we used an IL-4 dual reporter line, 4get/KN2 mice, in which eGFP expression marks cells expressing *Il4* mRNA (IL-4 competent) and surface huCD2 expression marks IL-4 producing Tfh cells in the mesenteric lymph nodes (mLN) draining the infection site^12^. Analysis of mLN from mice infected with either *Hpb*^SPF^ or *Hpb*^HA107^ larvae revealed a comparable induction of IL-4 competent and producing CD4^+^ T cells between groups (Figures 2C-E). As a complementary readout of Th2, we also assessed CD4^+^ T cell expression of GATA3, a transcription factor required for Th2 lineage commitment. Consistent with our eGFP results, *Hpb*^HA107^ elicited similar numbers of IL-4 producing GATA3^+^ Th2 cells as *Hpb*^SPF^ (Figures 2F & G).

The humoral component of the anti-*Hpb* immune response is dominated by the induction of the class-switched IgE and IgG1 antibodies, the latter of which is required for protective immunity to re-infection^13^. In line with the Tfh cell response, *Hpb*^HA107^ elicited comparable titres of serum IgE as well parasite-specific IgG1 to *Hpb*^SPF^ (Figures 2H & I). Overall, these data demonstrate that despite being raised in a limited bacterial monoculture, the effectively germ-free HA107-reared *Hpb* display robust infectivity and fecundity relative to SPF *Hpb* and are able to evoke a comparable immune response *in vivo*.

### Axenic helminth infection culminates in impaired parasite fitness

In order to validate that *Hpb*^HA107^ can maintain sterility in an *in vivo* setting, we infected germ-free mice and used several readouts to test microbial status. First, germ-free mice infected with *Hpb*^HA107^ (*Hpb*^HA107^ → GF) retained significantly larger ceca than germ-free mice infected with *Hpb*^SPF^ (*Hpb*^SPF^ → GF) (Figure 3A & B). Second, anaerobic and aerobic cultures using feces from mice at 14 days post-infection with *Hpb* ^HA107^ had no detectable bacterial growth in any conditions tested (see Methods for details) compared to controls. Third, DNA labeling of fecal samples with Sytox Green revealed no detectable bacteria in *Hpb*^HA107^ → GF mice, compared to abundant detection of bacteria in the *Hpb*^SPF^ → GF group (Figure 3C). Finally, a quantification of fecal 16S rDNA expression levels by qPCR also confirmed no appreciable increase in bacterial load in the feces of *Hpb*^HA107^ → GF mice after infection (Figure 3D). This result was in stark contrast to the over 1000-fold increase in 16S rDNA observed in the *Hpb*^SPF^ → GF group.

**Figure 3.**
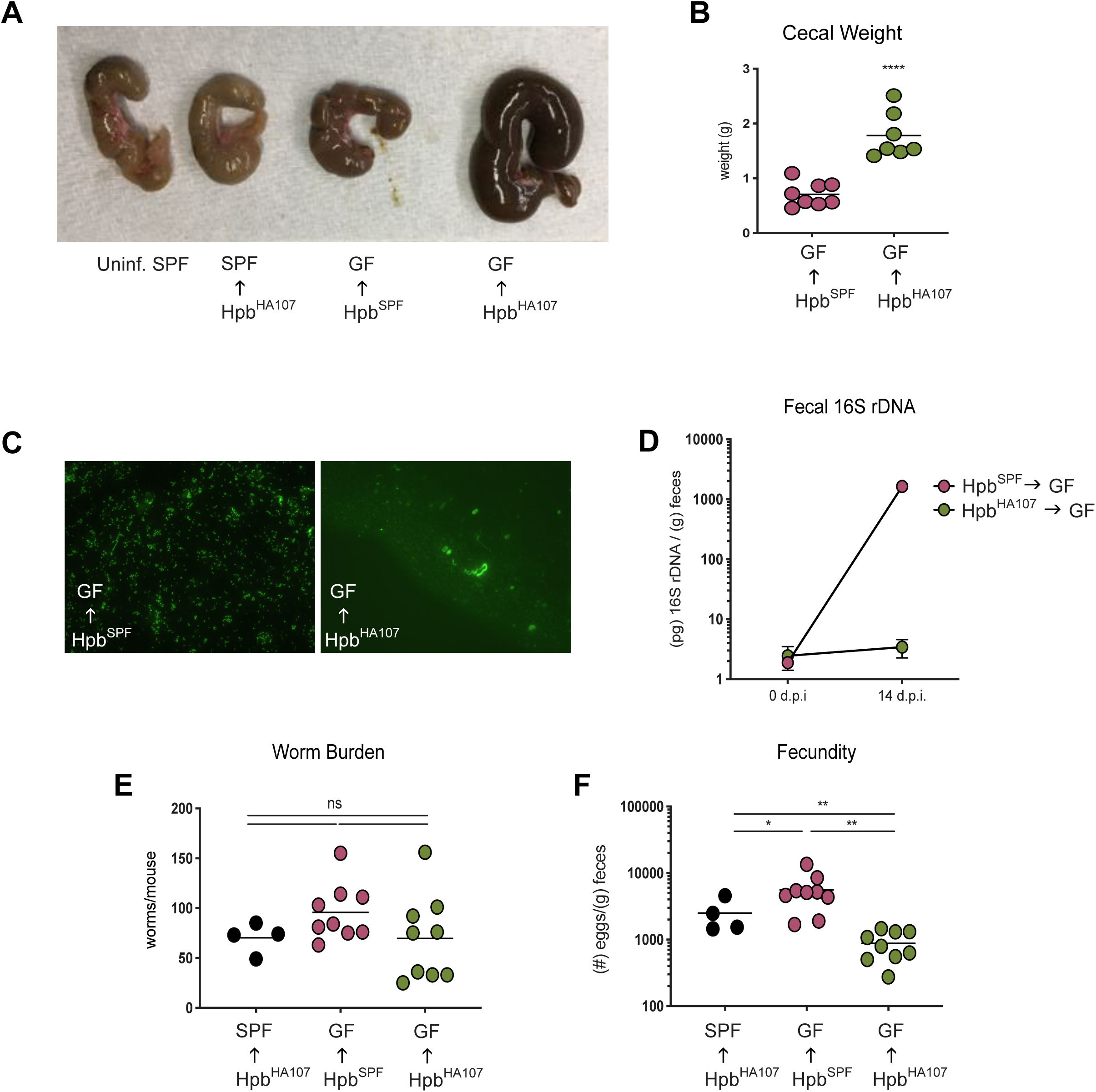
*Hpb*^HA107^ maintains host sterility and axenic helminth infection culminates in impaired parasite fitness. Germ-free (GF) C57BL/6 mice were infected with 200 HA107-reared or SPF L3 *Hpb* larvae (denoted *Hpb*^HA107^ → GF and *Hpb*^SPF^ → GF, respectively). (*Hpb*^HA107^ → SPF) indicates SPF C57BL/6 mice receiving 200 HA107-reared larvae. (Uninf. SPF) indicate uninfected SPF C57BL/6 mice. 14-day infected mice had their ceca excised and photographed (A), and cecal weights tabulated (B). Sytox Green staining of select cecal samples are shown (C). DNA was extracted from fecal samples and 16S rDNA (V6) levels before and after infection were quantified by qPCR (D). Adult worm burden (E) and fecundity (F) at 14 days post-infection are shown. Statistical analysis was performed using a t test with Welch’s correction. * indicates p < 0.05, ** indicates p < 0.01, *** indicates p < 0.001, ns = non-significant.

To begin exploring how the absence of a host microbiota affects the dynamics of *Hpb* infection, we examined the effects on worm burden and fecundity. No significant difference in worm burden was observed between axenic *Hpb* infection relative to SPF conditions at two weeks post-infection (despite a trending decrease) (Figure 3E). A significant reduction in worm fecundity, however, was observed in *Hpb*^HA107^ → GF mice in comparison to both *Hpb*^SPF^ → GF subjects and SPF mice receiving germ-free larvae (*Hpb*^HA107^ → SPF) (Figure 3F). Overall, these data demonstrate that *Hpb*^HA107^ effectively maintain germ-free status *in vivo* and that the microbiota promotes parasite fitness during helminth infection.

### Germ-free *Hpb* infection results in increased Th2 immunity and persistent intestinal granulomas

As enhanced Th2 immunity during primary *Hpb* infection has been shown to compromise parasite fitness^14^, we assessed key aspects of the anti-*Hpb* immune response to determine if any marked changes during infection accompanied the absence of a commensal microbiota. Analysis of mLN cells revealed a significant increase in GATA3+ Th2 cell frequency in germ-free mice infected with *Hpb*^HA107^ relative to all other groups, despite similar overall cell numbers (Figures 4A-C). This was accompanied by modestly elevated titres of IgE, as well as highly elevated parasite-specific IgG1 antibodies in the serum of these mice (Figures C & D).

**Figure 4.**
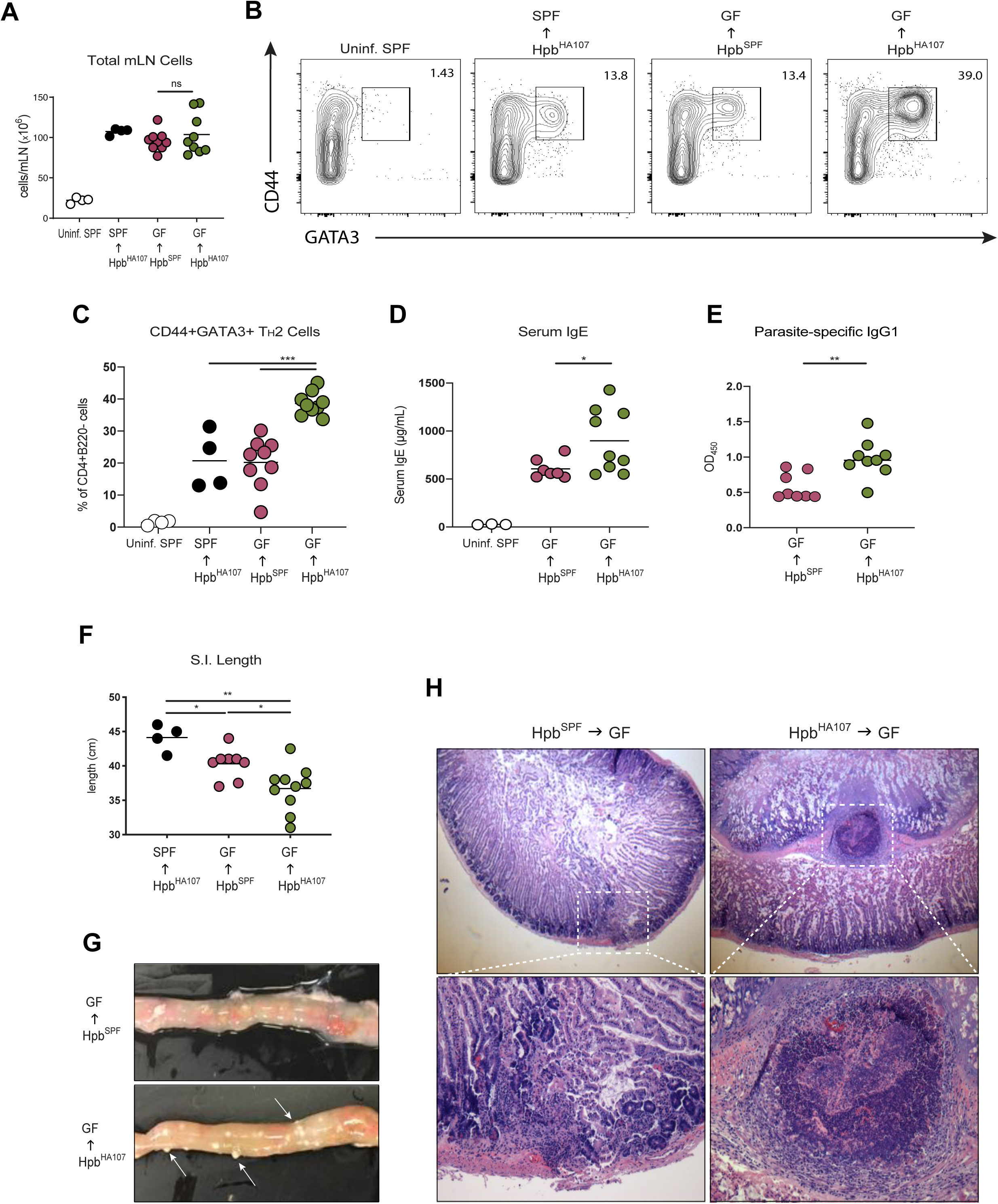
Germ-free helminth infection results in increased Th2 immunity and persistent intestinal granulomas. Germ-free (GF) C57BL/6 mice were infected with 200 HA107-reared or SPF L3 *Hpb* larvae (denoted *Hpb*^HA107^ → GF and *Hpb*^SPF^ → GF, respectively). (*Hpb*^HA107^ → SPF) indicates SPF C57BL/6 mice receiving 200 HA107-reared larvae. (Uninf. SPF) indicate uninfected SPF C57BL/6 mice. (A-C) 14 days post infection, mesenteric lymph nodes were taken and analyzed by flow cytometry. (A) Total mLN cell numbers, (B) representative plots and (C) frequencies of live CD4^+^B220^-^CD44^+^GATA3^+^ cells are shown. Serum was extracted and ELISAs for total IgE (D) and parasite-specific IgG1 (E) were performed. Murine small intestines were excised and measured (F). Photography, as well as H&E staining of duodenal sections are shown (G-H). Statistical analysis was performed using a t test with Welch’s correction. * indicates p < 0.05, ** indicates p < 0.01, *** indicates p < 0.001, ns = non-significant.

At the infection site, we observed a significant decrease in small intestinal length among *Hpb*^HA107^ → GF (Figure 4E). Furthermore, these mice displayed a uniform inability to resolve tissue granulomas (Figure 4F) – formations that typically arise during the tissue invasive stage of the parasite and normally resolve within 7-10 days post-infection^8, 15^. Indeed, at 14 days post-infection, all *Hpb*^HA107^ → GF subjects exhibited persistent granulomas that were not present in any other groups. Consistent with this result, histological examination of the small intestine revealed massive cellular infiltration within these granulomatous lesions that did not contain *Hpb* parasites (Figure 4G). Overall, these data demonstrate that an increased Th2 and type 2 humoral response, as well as dysregulated granuloma resolution, accompany impaired parasite fitness during axenic helminth infection.

## DISCUSSION

Our description of a detailed methodology for rearing effectively germ-free *Hpb* larvae expands on previous work^5^ and validates this model for *in vivo* studies. This technique will serve as an alternative to other methods of sterilizing *Hpb* such as antibiotic treatment of the L3 larvae prior to infection of germ-free mice^16^. Our methodology provides two improvements over such approaches. First, it avoids any possible fitness defects that could result from treating L3 *Hpb* with antibiotics or other sterilizing solutions. Second, our approach ensures that infectious larvae do not harbour any contaminating commensal microbes of their own that may not be accessible by exogenous treatment. Indeed, antibiotic treatment of L3 larvae likely only eliminates surface-dwelling microbes (since this is a non-feeding stage of parasite development). Since our methodology uses antibiotic sterilization at the egg stage (with the intention of avoiding fitness detriments of treating larvae) and then hatches the eggs into a pure monoculture of *E. coli* HA107, all the internal microbes of the developing L3 larvae should be unable to contaminate a germ-free host. Nevertheless, it remains possible that the limited microbiota of *Hpb*^HA107^ could lead to some minor fitness defects. However, our validation of *Hpb*^HA107^ infectivity and fecundity in SPF hosts demonstrates that their effects are negligible. Going forward, the use of germ-free *Hpb* will allow *in vivo* studies to examine microbiota-independent roles for helminths in immunomodulation and host fitness. Additionally, this model may serve as a valuable tool for refining *in vitro* studies examining helminth-intrinsic immune modulation.

Our observation that axenic helminth infection results in impaired parasite fecundity is consistent with previous reports^16, 17^. In addition to this reduction in fitness, we observed a non-significant, but trending decrease in worm burden in germ-free mice infected with *Hpb*^HA107^ at two weeks post-infection. Since decreased fecundity is often a precursor to worm expulsion, we speculate that the trending decrease in worm burden at two weeks post-infection may become significant at later time points of infection. Thus, the microbiota may impact fecundity, but also chronicity of primary infection. Although our data demonstrate that the commensal microbiota benefits the parasite during infection, it remains unclear whether this benefit is due to effects on the host via microbiota-mediated dampening of an anti-parasitic (type 2) immune response or by other unknown modifications of the intestinal microenvironment that supports the parasitic lifestyle. In support of the former idea, we observed an enhanced Th2 and parasite-specific humoral immune response, both of which are required for protective immunity to *Hpb* infection^13, 18^, under germ-free conditions. However, it remains to be determined whether other mechanisms including secretion of anti-helminth effector molecules by intestinal epithelial cells such as Relm-β and Pla2g1b, previously shown to compromise worm feeding and survival, may be increased in axenic conditions^19, 20^. As other enteric helminth species such as *Trichuris muris* rely on commensal microbes for their development, another non-mutually exclusive explanation is that the microbiota directly promotes *Hpb* fitness^21^. Indeed, it has been demonstrated by that *Hpb* elicits reproducible changes to the commensal gut microbiome^3, 22^. Whether these changes are directly mediated by the parasite and how they may specifically contribute to *Hpb* fitness is not clear.

The stark inability of germ-free mice to resolve helminth-induced granulomas confirms a previous report^17^ and demonstrates that the host also benefits from a commensal microbiota during *Hpb* infection. Based on our results, we speculate that persistent granulomas result from a dysregulated tissue repair process facilitated by an excessive Th2 response. We and others have recently described an early type 1 immune response, characterized by IFNγ-producing natural killer cells, in proximity to the *Hpb* granuloma that promotes disease tolerance and tissue regeneration during infection^15, 23^. Although it remains to be determined whether the microbiota is required for IFNγ induction in this context, this may be one mechanism by which the host exploits the microbiota to counter the type 2 immune response and promote granuloma resolution. Future studies using this model of axenic helminth infection will be of value in uncovering how the microbiota, or components of the microbiota, may limit the development of protective immunity against helminths.

## METHODS

### Animals and Infection

4get/KN2 and CD45.2 mice on a C57BL/6 background were bred and housed under specific-pathogen-free (SPF). Mice were used at 6-12 weeks of age and all experiments were approved by the McGill University Animal Care Committee. Germ-free mice were generated by a two-stage embryo transfer at McMaster University’s Axenic Gnotobiotic unit and shipped in germ-free containers (Taconic) to the gnotobiotic animal facility at the Institut Universitaire de Cardiologie et de Pneumologie de Québec (IUCPQ), Université Laval. After seven days of acclimation, germ-free mice were infected by gavage with 200 L3 *Hpb* larvae (either SPF- or HA107-grown) and harvested at the indicated time points.

### Generation of *Hpb*^SPF^ Larvae

Infective L3 stage *Hpb* larvae were generated through an adapted methodology of a previous publication^9^. In short, the feces of *Hpb*-infected Balb/c mice were taken and spread onto moist Whatman filter paper in a 10cm petri dish and generously doused with sterile water. Over the course of several days, the fecal cultures were watered and infective larvae were collected and stored at 4°C.

### Generation of *Hpb*^HA107^ Larvae

4get/KN2 mice at least 6 weeks of age were gavaged with 400 (fecal-grown) L3 *Hpb*. Between 2 and 4 weeks post-infection, mice were sacrificed and the first 10 cm of their duodenum were excised. Under a laminar flow hood, the murine small intestines were opened longitudinally and, using sterile forceps, adult *Hpb* worms were extracted and placed into RPMI containing 0.1mg/mL metronidazole, 0.1mg/mL ampicillin, 0.05mg/ml vancomycin, 0.1mg/mL neomycin, 100 IU/mL penicillin, 0.1mg/mL streptomycin, 1mg/mL gentamycin, 2.5μg/mL Amphotericin B, and 1% D-Glucose. Care was taken to avoid transferring large quantities of mucus or luminal contents. After extraction, the adult worms were transferred into a 50mL tube, washed with sterile PBS, and allowed to settle by gravity. The supernatant was removed and the worms were washed at least 8 more times with a full 50mL of sterile PBS, until the supernatant was clear. After the final wash, the adult worms were transferred into a 10cm Petri Dish containing at least 25mL of RPMI containing 0.1mg/mL metronidazole, 0.1mg/mL ampicillin, 0.05mg/ml vancomycin, 0.1mg/mL neomycin, 100IU/mL penicillin, 0.1mg/mL streptomycin, 1mg/mL gentamycin, 2.5μg/mL Amphotericin B, 1% D-Glucose, and 10% FBS, and left to incubate at 37°C. After 18 hours, the worm-containing media was strained through a 70μm filter into a fresh 50mL tube under a laminar flow hood. The flow-through was centrifuged at 2200rpm for 4 minutes, and the supernatant was pipetted off the pellet of *Hpb* eggs. The pellet was re-suspended in 5 mL of sterile ddH_2_O and transferred to a 15mL tube wherein it was washed 3 more times with sterile H2O (performing spin steps at 700g for 2 minutes). After the third wash, the eggs were re-suspended in 14mL of RPMI containing 0.2mg/mL metronidazole, 0.2mg/mL ampicillin, 0.1mg/ml vancomycin, 0.2mg/mL neomycin, 200IU/mL penicillin, 0.2mg/mL streptomycin, 2mg/mL gentamycin, and 5μg/mL Amphotericin B, and left to incubate at 4°C for 24 hours. After incubation, the eggs were washed a minimum of 6 times in sterile ddH_2_O, spinning at 700g for 2 minutes. The sterility of the eggs was confirmed by plating on LB and YPD media. The eggs were re-suspended in ∼10mL of ddH_2_O, counted, centrifuged, and re-suspended in the appropriate volume of water to achieve roughly 20 eggs/μl. In sterile Eppendorf tubes, roughly 5000 *Hpb* eggs were combined with 100μl of pure *E. coli* HA107 overnight culture (OD600 = 0.650 to 1.00). If necessary, dilutions of HA107 were performed using appropriately supplemented LB media^10^. The egg-bacteria mixture was plated onto Nematode Growth Media (NGM), and the plates were gently tilted in an orbital manner to spread the mixture into a circle (leaving a gap between the liquid and edge of the dish). NGM was made by combining 3.0g NaCl, 2.5g peptone, 17g agar in 975mL ddH_2_O. After autoclave sterilization and cooling, 1M each of CaCl_2_, MgSO_4_, and potassium phosphate buffer were added, along with 5mg/mL cholesterol (dissolved in ethanol). The monoculture/*Hpb*-seeded plates were placed into sterile plastic bags, sealed, and left at room temperature in the dark. After 2-3 days, plates were lightly watered with 1mL of sterile ddH_2_O and re-bagged. After a total of 5 days, the L3 larvae were harvested by gently pipetting sterile water onto the surface of the agar. Larvae were immediately placed into 15mL tubes and washed 3-4 times with sterile water (kept on ice during collection). Preliminary L3 sterility was confirmed by plating on LB and YPD media and larvae were stored at 4°C. Larvae were used no more than one week after harvest. Prior to infection, worms were washed again 2-3 times with water, and 50uM EDTA was added to the worm suspensions to prevent adherence to the tubes.

### *Hpb* egg quantification

At least 0.2g of feces was collected and vortexed in a 0.75-1mL saturated NaCl solution. Suspensions were incubated at room temperature for 24 hours, followed by at least 6 days at 4°C. Then, the top 2 layers (clear and milky) were extracted and centrifuged at 4000rpm for 4 minutes. Supernatants were discarded and egg pellets were re-suspended in ddH_2_O for counting.

### Fecal DNA extraction and qPCR

Fecal DNA was extracted using the QIAamp Fast DNA Stool Mini Kit (QIAGEN) according to the manufacturer’s instructions. qPCR for the V6 hypervariable region of the 16S rRNA gene segment was performed using the following primers: aggattagataccctggta (forward), cttcacgagctgacgac (reverse).

### Microbial detection assays

In experiments using germ-free mice, fresh cecal contents were collected and stained with Sytox Green (Thermofisher) for detection of bacterial DNA. In addition, fecal samples were obtained from the same mice at day 14 post-infection with *Hpb* andincubated in brain heart infusion broth (BHI) for 7 days in three conditions (37°C and 25°C aerobic and 30°C anaerobic). Positive samples were observed by microscopy, Gram staining and further plated on Brain heart infusion Agar (BHIA) for colony-forming unit isolation and their DNA extraction. All samples from germ-free mice infected with *Hpb*^HA107^ tested negative for bacterial contamination.

### Flow Cytometry

mLN were dissected from uninfected or infected mice and filtered through a 70μm strainer into PBS containing 10μM HEPES and 10% FBS. Live cells were counted with a haemocytometer by trypan blue exclusion. Single cell suspensions were stained with fixable viability dye (EBioscience) in PBS. Following viability staining, cells were incubated with anti-Fc receptor (clone 2.4G2, BD Biosciences) for 7-12 minutes. Cells were immediately surface stained for 30 minutes on ice. The following antibodies were used: CD4-PerCP (Ebiosciences: GK1.5), CD4-BUV395 (BD: GK1.5), CD4-APC-Cy7 (Ebiosciences: GK1.5), B220-PE-Cy7(Ebiosciences: RA3-62B), B220-FITC (invitrogen: RA3-6B2), CD44-BV421 (BD: IM7), CD44-PE (Ebiosciences: IM7), CD62L-APC (EBiosciences: MEL-14), huCD2-PE (Ebiosciences: RPA-2.10), PD1-PE-Cy7 (EBiosciences: J43), CXCR5-biotin (Ebioscienes: SPRCL5), SA-APC (Ebiosciences). In experiments where intracellular staining was performed, cells were fixed using an Ebioscience Transcription Factor Staining Buffer Set. The following antibodies were used for intracellular staining: GATA3-PerCP-eF710 (invitrogen: TWAJ), FoxP3-AF700 (Ebiosciences: FJK-16s), Data were acquired with a FACSCanto II II or LSR Fortessa (BD Biosciences). Results were analyzed using FlowJo software.

### ELISAs

Total serum IgE was quantified using a Mouse IgE ELISA Ready-SET-Go! kit (eBio/Affymetrix). HES-specific IgG1 was quantified using a Mouse IgG1 ELISA Ready-SET-Go! kit (eBio/Affymetrix). HES was prepared as previously described^9^ and a 96-well flat-bottom plate was coated with 1µg/mL and incubated at 4°C overnight. Rat anti-mouse IgG1-Biotin (SB77E) and Streptaviding-HRP (Southern Biotech) were used as detection and secondary antibodies, respectively.

### Larval Growth Images

All larval growth photos were taken at 10X magnification using an Olympus BX50 microscope.

### Histology

Duodenal samples were formalin-fixed, paraffin-embedded, sectioned and stained with hematoxylin and eosin by the MUHC-RI Histology Core.

### Statistical Analysis

Data were analyzed by a student’s t test using the GraphPad Prism program (versions 7 & 8). A p value < 0.05 was considered significant.

## ACKNOWLEDGMENTS

We would like to thank Dr. Andrew J. Macpherson for graciously providing the *E. coli* HA107, Dr. Victoria Zismanov for help with flow cytometry and Dr. Christian E. Rocheleau and Sarah Gagnon for their experimental support in the development of the model. We would further like to acknowledge members of the King Laboratory for their technical input. E.F.V. holds a Canada Research Chair in Nutrition, Inflammation and Microbiota. G.A.R. is a recipient of a McGill Science Undergraduate Research Award (SURA). I.L.K. holds a Canada Research Chair in Barrier Immunity which supported the completion of this work.

## AUTHOR CONTRIBUTIONS

G.A.R. designed and performed experiments, analyzed data, and wrote the manuscript. C.F. performed the helminth infections in the gnotobiotic animal facility at the IUCPQ, Université Laval. E.F.V. provided experimental support and intellectual input. All authors critically reviewed the manuscript. I.L.K designed the project and wrote the manuscript.

